# Survival of prey growing through gape-limited predators

**DOI:** 10.1101/571356

**Authors:** James J. Anderson

## Abstract

Juvenile to adult survival of fish is modeled by the rate at which prey progressively escape the size distribution of gape-limited predators through growth. The model characterizes adult survival as a function of the mean and standard deviation of the predator population gape sizes, the ratio of mortality and growth rates and a compensatory growth factor. The model fits the survival of adult returns of Chinook salmon and reveals that a 25% increase in either the initial size at ocean entrance or the growth to mortality rate over the first few months of ocean residence can increase adult survival by a factor of 2 to 3. Additionally, the model proposes a rigorous mechanism through which the size distribution of predators determines the effects of juvenile growth on adult survival. Finally, possible contributions of the model framework to fisheries management and predator-prey theory are noted.

## 1. Introduction

In many species, mortality exhibits an age-declining rate that results in the population survival being strongly influenced by early life predation [1]. The effect of age-specific mortality is of particular importance in fishes where predation is the dominant form of mortality [2, 3]. Susceptibility to predation typically depends on the size of the prey relative to the gape size of the predators [4, 5]. The mechanism, denoted as size-selective mortality [3, 6–10], involves the interplay of the predator and prey sizes and prey growth rate, which leads to larger fish surviving better than small ones to adult stages [11, 12].

To illustrate the dramatic contrast of mortality with size consider the marine survival of spring Chinook salmon (*Oncorhynchus tshawytscha*) from Snake River hatcheries in the Columbia River Basin of the Pacific Northwest, USA. Using data extracted from [11], salmon enter the ocean at lengths of 10-15 cm and return as adults at lengths of up to a meter. The ratio of the number of adult in a cohort returning to the river to spawn to the number of the juveniles in the cohort that entered the ocean, denoted the smolt-to-adult-ratio (SAR), is on the order of 1% and the average mortality rate over the ocean residence is ~0.00002/d. In contrast, the juvenile mortality rate in the first few weeks in the ocean is ~ 0.1/d [13]. This 10^4^ differential in rate measures suggests the mechanism involves size-selective mortality [14, 15], which also is well documented in other fishes [3, 6–10]. However, the process is complex. Salmon SAR characteristically increases with smolt length at marine entrance [11, 12] but not in all populations [16–18]. In some Chinook salmon populations, size-selective mortality was detected during early ocean residence [14, 15, 19], but only after the first 30 days of marine residence in others [8] and in freshwater stages in yet other populations [6]. Additionally, juvenile size and adult return relationships can be different for natural and hatchery reared juveniles with smaller wild fish survival exceeding the survival of hatchery counterparts of the same size [17].

While models of size-selective mortality are abundant in the literature, arguably they do not provide clear explanations of these complexities. The classic Sinko-Streifer model [20] allow analytical solutions of survival in terms of the interaction of aggregated measures of animal physiology with age and size. However, the solutions are of limited use because the growth and mortality processes are highly simplified. In particular, such models typically do not incorporate details of the predator size or effects of size on prey growth. Individual based models (IBM) overcome these limitations by expressing the properties of the predators and prey and rules for their interactions [3, 21]. While IBMs express more realism, it comes at the expense of a high number of variables and computational limitations that make applications of IBMs to management of ecosystems and commercial fisheries difficult.

This paper presents a model of intermediate but tractable complexity. It characterizes size-dependent predator-prey interactions by integrating predator-prey size differential, predator abundance and prey growth properties in a Lagrangian framework. In essence, the age-specific mortality rate is cast in terms of the duration of time required for the prey to grow larger than the gape sizes of predators.

## 2. Materials and methods

### (a) Model

The model is based on the idea that prey susceptibility to predation decreases as prey progressively grow larger. The prey survival is expressed

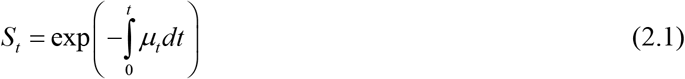

where *μ_t_* is the mortality rate as a function of prey age *t* where *t* = 0 is some reference age when the prey enters the predator field. To incorporate size dependency in the mortality rate, assume predator-prey encounters have a Poisson distribution with mean frequency *λ* and for an encounter to result in prey mortality the size of the prey *l* must be less than the size of the predator’s gape, *Z*, i.e. *Z* > *l*. Applying a formulation developed for vitality-based human survival [22], the mortality rate is 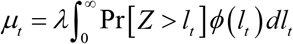 where Pr[*Z* > *l_t_*] is the probability of the predator being larger than prey of size *l_t_* at *t* and *ϕ*(*l_t_*) is the distribution of prey lengths at time *t*, which depends on both the prey growth rate and the preferential elimination of smaller prey through predation. There is no closed form for *ϕ*(*l_t_*) so applying a discretized approach [23] the mortality rate is tracked for size class increments, i, defined by the size of fish at *t* = 0. The class-specific mortality rate is then approximated as *μ_i,t_* = *λ* Pr[*Z* > *l_i,t_* and size class-specific survival is

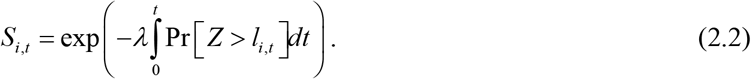

To express the probability that a predator gape size is greater than the prey size represent the predator size distribution by a truncated cumulative normal distribution *F*(*l, m, s, a, b*) where l is the prey length, *m* and *s* are the mean and standard deviation of the predator gape distribution, and the truncation interval is *a* = 0 to *b* = ∞. The probability of the predator exceeding the prey size is 1 − *F*, which yields

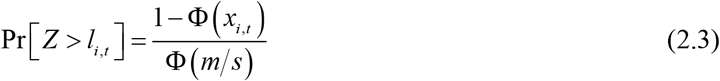

where Φ is the standardized cumulative normal distribution and the prey size normalized to the predator gape distribution is

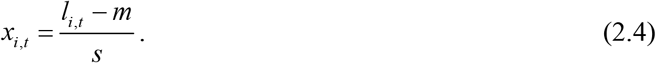

The mortality rate per size class i is then

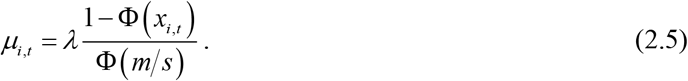

Figure 1 illustrates Pr where the prey size *l* = 0 corresponds to *x* = −*m/s* and prey size *x*_*i*,0_ = 0 corresponds to mean predator gape size.

**Figure 1.**
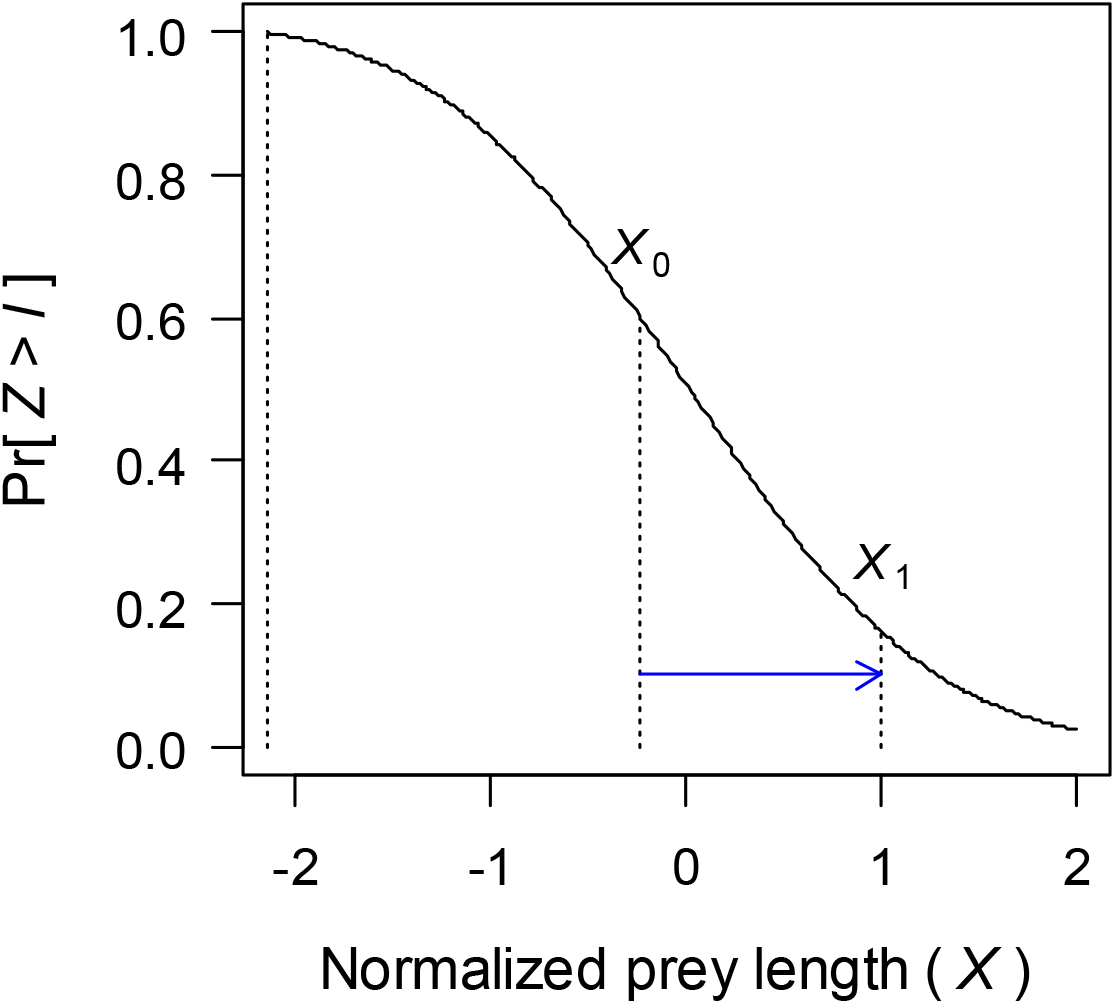
Normalized relationship of prey length to the predator gape size distribution according to equation (2.3) using *m* = 160 mm and *s* = 75 mm. Fish enters predator field at size *X*_0_. Fish growth to escape predator field depicted by blue arrow, where at reaching *X*_1_ the prey is larger than the gape of 84% of the predators. Normalized predator mean gape size is 0.

To include effects of age in the model assume linear prey growth as *l_i,t_* = *l*_*i*,0_ + *g_i_t* where *g_i_* is the growth rate in the class *i* and *l*_*i*,0_ is the prey length when it enters the predator field at *t* = 0. Then the normalized prey size as a function of time is

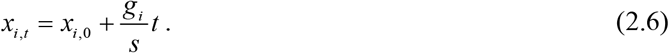

Equation (2.6) is similar to the early life growth in a Von Bertalanffy growth equation [24]. While at older ages the linear equation overestimates size and correspondingly underestimates mortality, the bias has little effect in the model because the majority of size-dependent mortality occurs in early age. Introducing equations (2.6) and (2.3) into (2.2) the log survival for size class *i* is

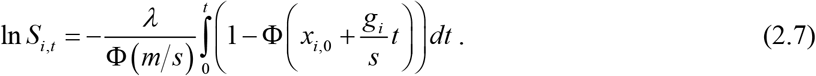

From equation (2.6) 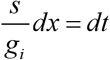 and expressing limits of integration in terms of *x*_*i*,0_ and *x_i,t_* then the integration of equation (2.7) gives the survival as a function of initial size and size at age *t* as

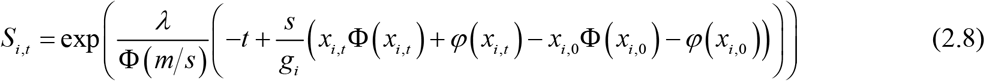

where *φ* is a standard normalized distribution. Noting as *t* → ∞ that *x_i,t_* − ∞, Φ(*x_i,t_*) → 1 *φ*(*x_i,t_*) → 0 and Φ(−x) = 1 − Φ(*x*) then equation (2.8) approximates the adult survival for size class *i* as

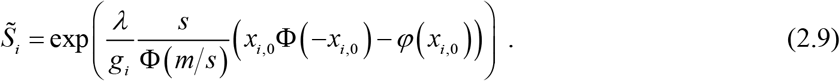

To express equations (2.8) and (2.9) in terms of the population average length and growth rate represent size-class growth rate relative to the average growth rate by a logistic curve

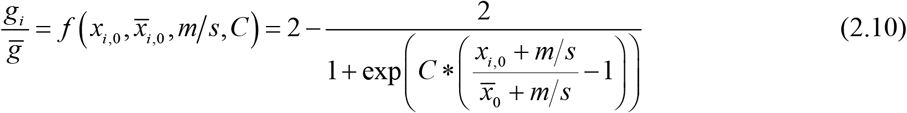

where *x*_*i*,0_ and 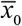 are the normalized prey size for class i and the mean size class of the population respectively at time *t* = 0, *ḡ* is the mean growth rate and *C* is a coefficient characterizing the relationship of size class and growth. For *C* = 0 growth rates are equal for all size classes. For *C* > 0 the growth rate is greater than *ḡ* for size classes larger than the mean, which suggests the possibility that differences in size at *t* = 0 was the result of different previous growth rates and the pattern is maintained for *t* > 0. Correspondingly, *C* < 0 suggests growth rates of the smaller size classes are greater than the mean population growth rate for *t* > 0. This condition could indicate compensatory growth in which prey with slower growth for *t* < 0 as a result of past diet restriction exhibit catch up growth when the diet restriction is released for *t* > 0 [25, 26].

Note from equation (2.4) that the fraction 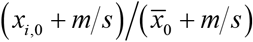 is simply 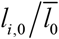 where 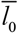 the mean length of the population at *t* = 0. Also note from equation (2.10) that 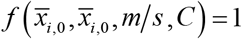.

Substituting equation (2.10) into (2.8) the gape-limited survival with age is

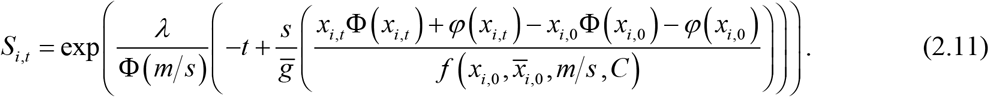

Mortality may also occur independent of the prey length and exposure time; for example when juvenile [27] or adult [28] salmon pass a gauntlet of predators or physical obstructions during their respective downstream and upstream river migrations. This mortality can be incorporated as a constant harvest rate *H* and the final equation for adult survival is

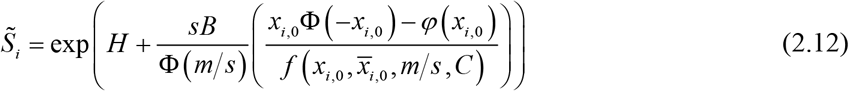

where

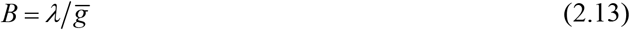

and the inverse 1/*B* is an index of the increase in fish length between predator encounters. Noting *s* is a measure of the thickness of the predator field in units of gape size then *N* = *sB* is an index of the number of predator encounters that occur in escaping gape-limited predation.

Finally, the ratio of the mean growth rate to the mean predation rate from gape-limited predation is

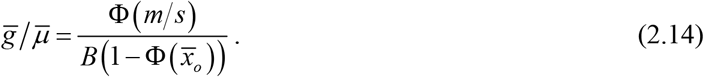

### (b) Example

To demonstrate the model, consider the ocean survival of spring Chinook salmon from Snake River hatcheries using data extracted from [11]. Juvenile smolts were collected, measured for length and tagged with passive integrated transponders at Lower Granite Dam on the Snake River. Data from two years are used in the example; fish tagged in 2008 were transported around the eight dams of the Snake-Columbia River hydrosystem in barges and fish tagged in 2009 were released back into the river. For both passage routes, the SAR from juveniles passing Bonneville Dam, the last dam in the system, to returning adults were calculated for 3 mm increments of size classes constructed by grouping adults according to their juvenile length. These data were selected because they represent the range of SAR observed for Snake River salmon.

The gape distribution of the marine predators was estimated from the size distributions of piscivore and avian predators residing in the Columbia River plume and the Washington coast. Pacific hake and jack mackerel, are major piscivore predators and through stomach content their prey length have been observed to cover the range between 50 and 250 mm with a median length of about 150 mm [29]. Seabirds also are important predators of juvenile salmon in the Columbia River plume and coastal environment. Aucklets foraging off Vancouver Island select salmonids at ~120 mm [30]. These estimates cover the range or gape size the juvenile salmon are expected to encounter during early marine residence.

Returning adult Columbia River salmon are also susceptible to pinniped predation (~4.5%) in the bypass system of Bonneville Dam, the first dam they encounter in the hydrosystem [28]. Adults caught in human harvest are accounted for in the SAR estimates and thus harvest is not included in the model.

The ratio of growth rate to the mortality rate during fish early marine residence is a fundamental measure that constrains the model parameters according to equation (2.14). For Snake River spring Chinook salmon, these rates have been estimated over the first few months of marine residence. Growth rates were obtained from the growth rings of otoliths removed from Snake River spring Chinook captured off the Washington coast. Estimate of growth over the first month in the ocean ranges from 0.5 to 1 mm/d [31]. Using data of acoustically tagged juvenile spring Chinook salmon transiting through the Columbia River plume [13], the mortality rate was estimated at *μ* ~ 0.08/d. The estimate is based on a log-linear regression of survival vs plume residence time and includes an intercept to account for experimental effects including body tag burden and capture inefficiency of the acoustic detection array. The resulting growth to mortality ratio is *ḡ/μ* ~ 11 ± 1.9

### (c) Estimating parameters

In principle, the parameters of equation (2.12) can be estimated with nonlinear estimation routines such as the nls package in the R statistical software [32]. However, a range of *m, s* sets produce essentially the same goodness of fit of SAR vs juvenile length and so a unique best-fit estimate of the parameters cannot be obtained from such regressions alone. Fortunately, the predator distribution can be estimated independently and equally well-fitting *m, s* sets produce different *ḡ/μ* ratios and *C*. In turn, the *ḡ/μ* ratio can be estimated from field studies of growth and mortality. The harvest parameter *H*, representing mortality not associated with time of exposure or prey length, can be problematic when included as a free parameter in the regression. However, *H* again can be estimated independently. For the Chinook salmon example, the effect of *H* was explored by comparing results with *H* = 0, corresponding to no harvest mortality, and *H* = −0.047, corresponding to the pinniped mortality reported by [28]. The resulting estimates of *B* and *C* differed by less than 2% so for demonstration purposes *H* = 0.

## 3. Results

Figure 2 illustrates the effect of the predator gape distribution on the model fit standard error and parameters. The contours were generated using the nls package to fit the 2008 and 2009 data for 1/3 mm increments of *m* and *s*. For both years the fitting standard errors decrease with increasing m and s resulting in diagonal isopleths. Standard error isopleths depicted as dashed lines in figure 2 define transitions in the model goodness of fit. Above the transitions *m*, *s* sets produce equally good fits and sets below the transitions fit poorly. The transition standard error isopleths are 5.5 x 10^-3^ for 2008 and 1.0 x 10^-3^ for 2009. Correspondingly, the *ḡ/μ* isopleths exhibit a similar diagonal pattern but with the ratio linearly increasing with increasing *m* and *s*. The *C* isopleths also exhibit a diagonal pattern with a similar transition. Left of the −2.5 isopleth *C* is essentially constant for all *m*, *s* sets and *C* increases above the transition isopleth. Finally, the isopleths of B exhibit a diagonal pattern exponentially increasing with decreasing *m* and *s*. The examples of *m*, *s* sets (a, b, c, d) cover the region of good model fits to the SAR vs length data. The parameter values for these sets are given in table 1. The model fits to the SAR vs length data for 2008 and 2009 are illustrated for the four *m*, *s* sets in figure 3.

**Figure 2.**
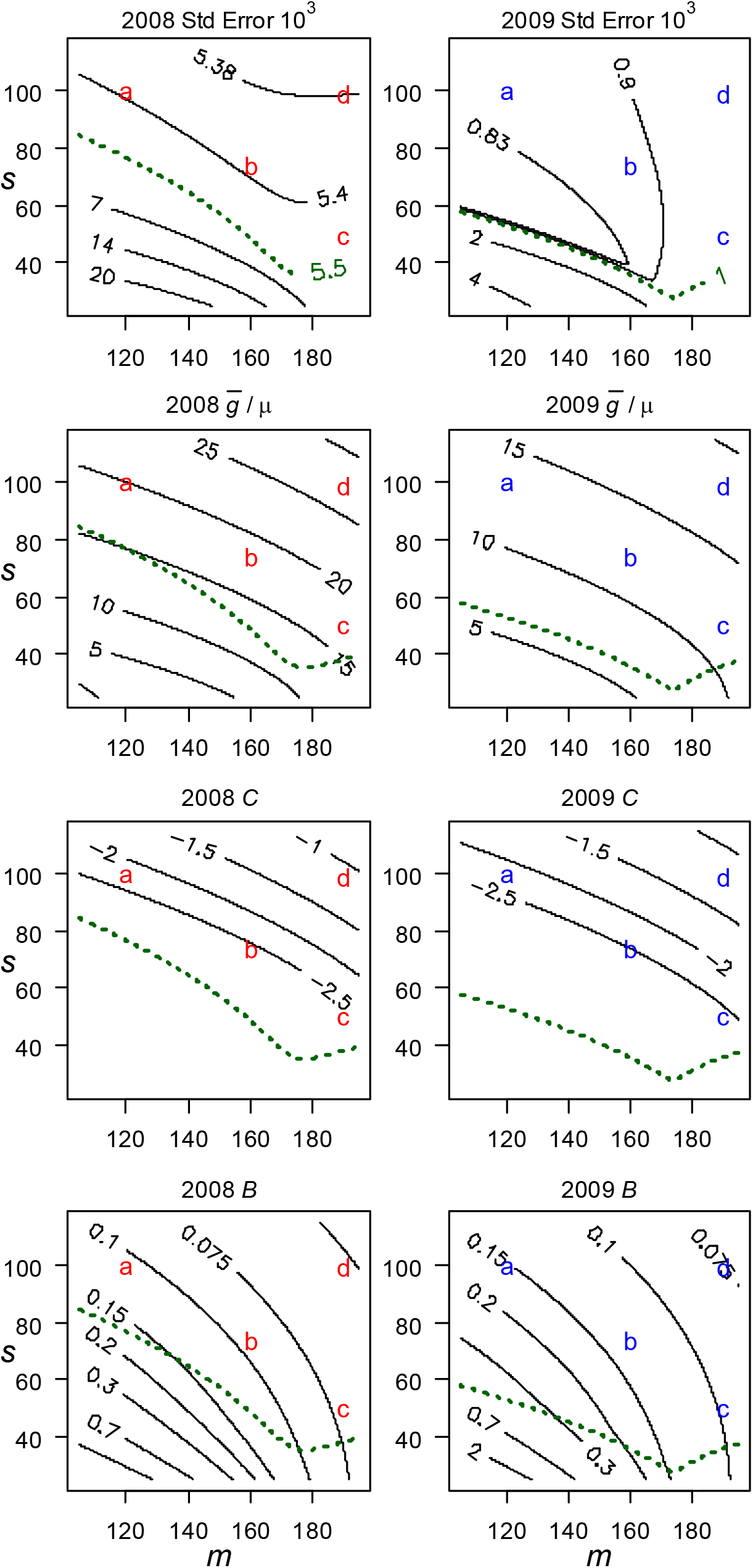
Effects of predator distribution parameters (*m, s*) on nls fits of equation (2.12) parameters for Snake River spring Chinook salmon in outmigration years 2008 and 2009. Points (a, b, c, d) correspond to *m, s* sets used in table 1. Dashed lines depict threshold standard error isopleths of s, m values with good model fits.

**Table 1.** Parameters for Snake River spring Chinook salmon migration year 2008 and 2009 with 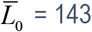 and 135 mm respectively. See text for details.

| Year | Set | $m$<br>mm | $s$<br>mm | std err<br>$10^3$ | $\bar{g}/\mu$<br>mm | $C$<br>- | $B$<br>1/mm | $\lambda$<br>1/d | $\bar{g}$<br>mm/d |
| --- | --- | --- | --- | --- | --- | --- | --- | --- | --- |
| 2008 | a | 120 | 100 | 5.40 | 19.97 | -2.22 | 0.11 | 0.11 | 1 |
| 2008 | b | 160 | 75 | 5.39 | 18.40 | -2.54 | 0.09 | 0.09 | 1 |
| 2008 | c | 190 | 50 | 5.43 | 17.20 | -2.80 | 0.07 | 0.07 | 1 |
| 2008 | d | 190 | 100 | 5.38 | 27.46 | -1.09 | 0.05 | 0.05 | 1 |
| 2009 | a | 160 | 75 | 0.89 | 12.35 | -2.45 | 0.13 | 0.11 | 0.71 |
| 2009 | b | 190 | 50 | 0.95 | 11.91 | -2.65 | 0.10 | 0.09 | 0.72 |
| 2009 | c | 190 | 100 | 0.91 | 18.22 | -1.18 | 0.08 | 0.07 | 0.72 |
| 2009 | d | 190 | 100 | 0.88 | 18.2 | -1.2 | 0.08 | 0.05 | 0.69 |

**Figure 3.**
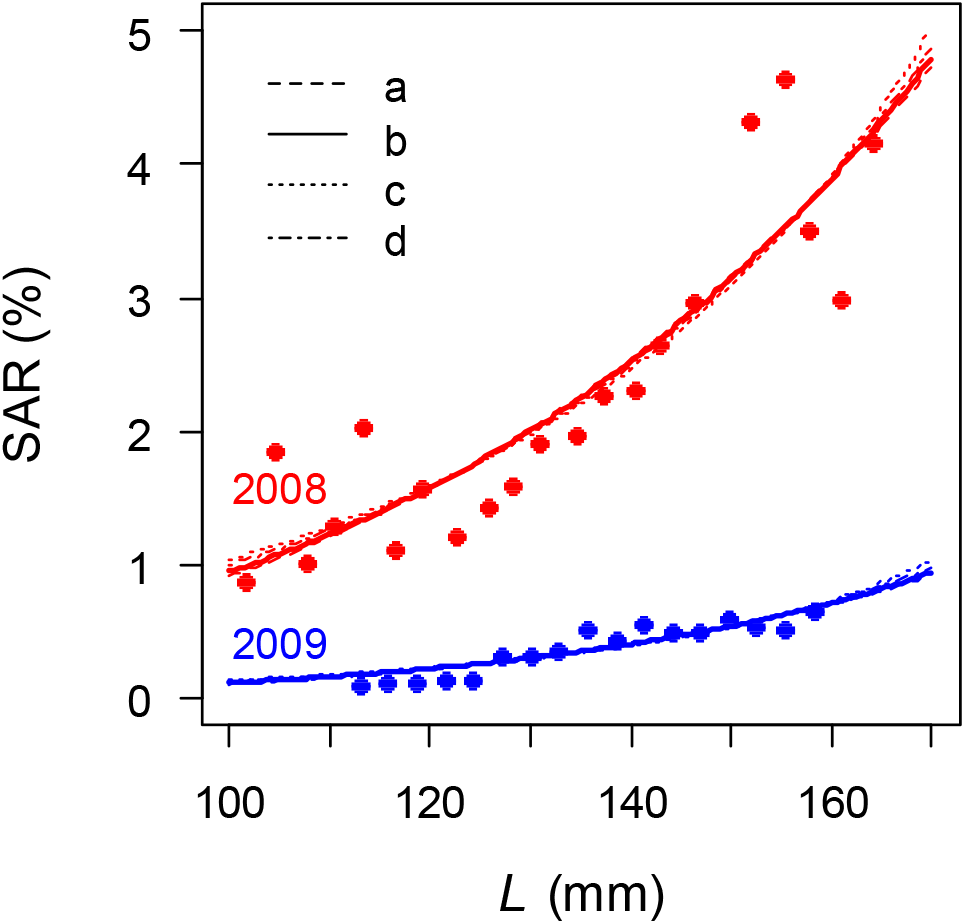
SAR vs length at juvenile tagging of Snake River spring Chinook salmon for 2008 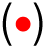 and 2009 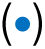. Lines depict equation (2.12) fits to data using predator gape distribution sets a, b, c, d from table 1.

Figure 4 illustrates modeled length, mortality rate, and ocean survival of the Chinook salmon for 2008 and 2009 using set b parameters. Additionally, figure 1 illustrates the predator gape distribution for set b. The patterns in figure 4 are based on a growth rate *ḡ* = 1 mm/d for 2008, which results in a predator encounter rate of 0.9/d. Assuming the same encounter rate for 2009, from equation (2.13) the 2009 growth rate is *ḡ* = 0.72 mm/d. The mortality rate declines more slowly for 2009 resulting in greater reduction in survival over the first 200 days of ocean residence compared to 2008. At 200 days, the 2009 fish length of 235 mm is larger than 84% of the predator gapes. Sets a, c, and d exhibit similar patterns.

**Figure 4.**
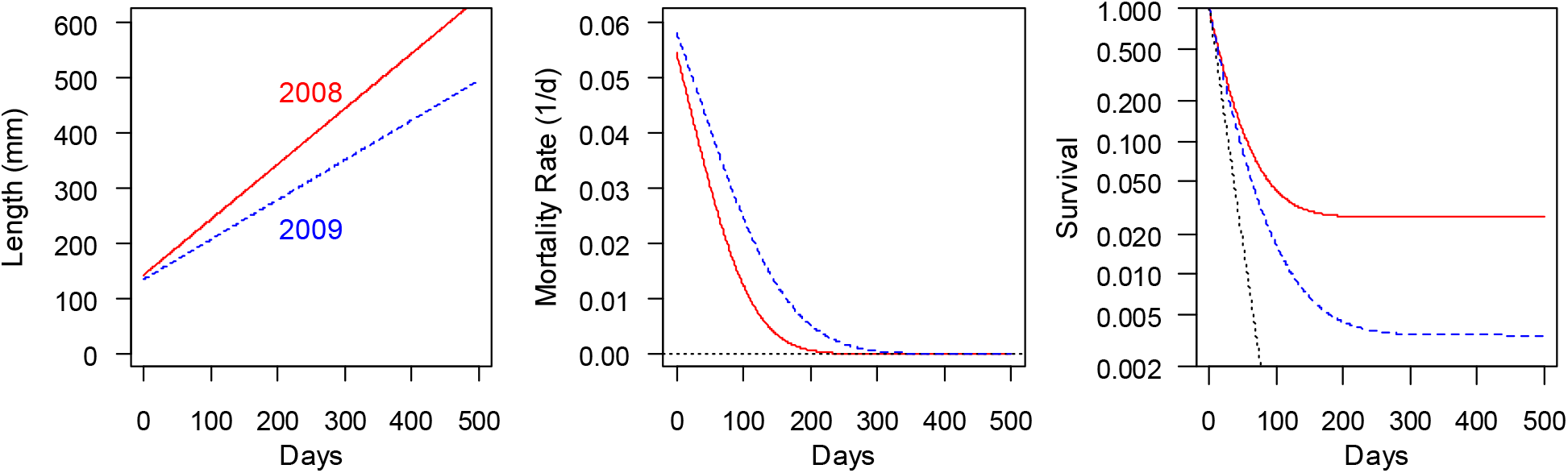
Distributions of growth, equation (2.6), mortality rate, equation (2.5), and survival, equation (2.11), for the 2008 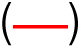 and 2009 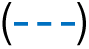 groups using the set b parameters in table 1. Dotted line (…..) in the survival figure corresponds to ln *S* = −0.08*d* using data from [13].

The effect of fish size on growth rate can be significant as revealed in figure 5 in which *C* between 2 and 3 results in smaller fish growth being 40% higher than the mean cohort growth rate. This suggest compensatory growth contributes to the SAR vs length pattern. It is noteworthy that *C* was negative, which is consistent with classical growth theories e.g. von Bertalanffy growth model [24] in which growth rate declines with increasing size.

**Figure 5.**
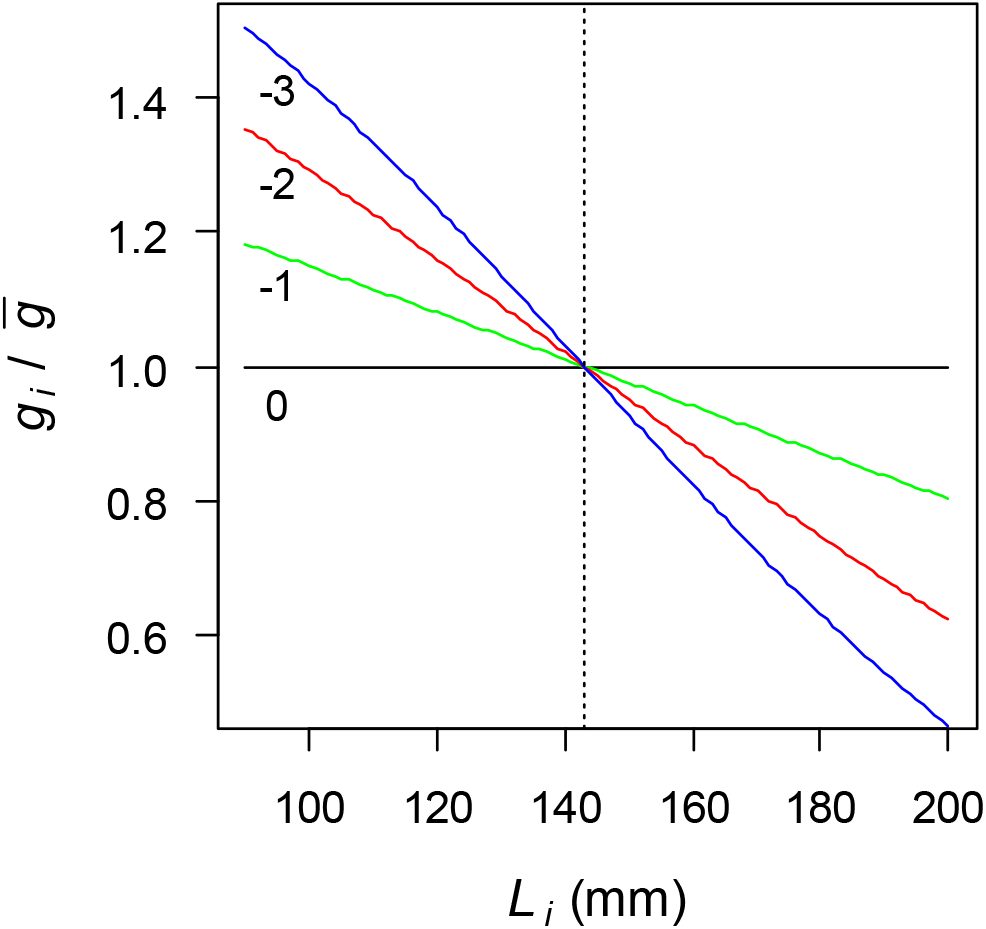
Effect of fish length on relative growth rate, equation (2.10) for values of *C*.

For a given predator gape distribution, the initial prey length and ratio of prey growth to mortality rates are main factors determining adult survival. Figure 6 illustrates these effects for predator distribution sets a-d. In each figure the growth to mortality rate ratio and initial length are individually increased by 25% from a base of *ḡ/μ* = 15.4 and 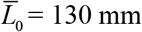 mm. While the predator size distribution affects the base SAR in each case, the relative increases are similar varying by factors between 1.8 and 2.5. A similar magnitude of effect was reported by [31] in which the SAR increased by greater than a factor of 3 for a 40% increase in growth rate.

**Figure 6.**
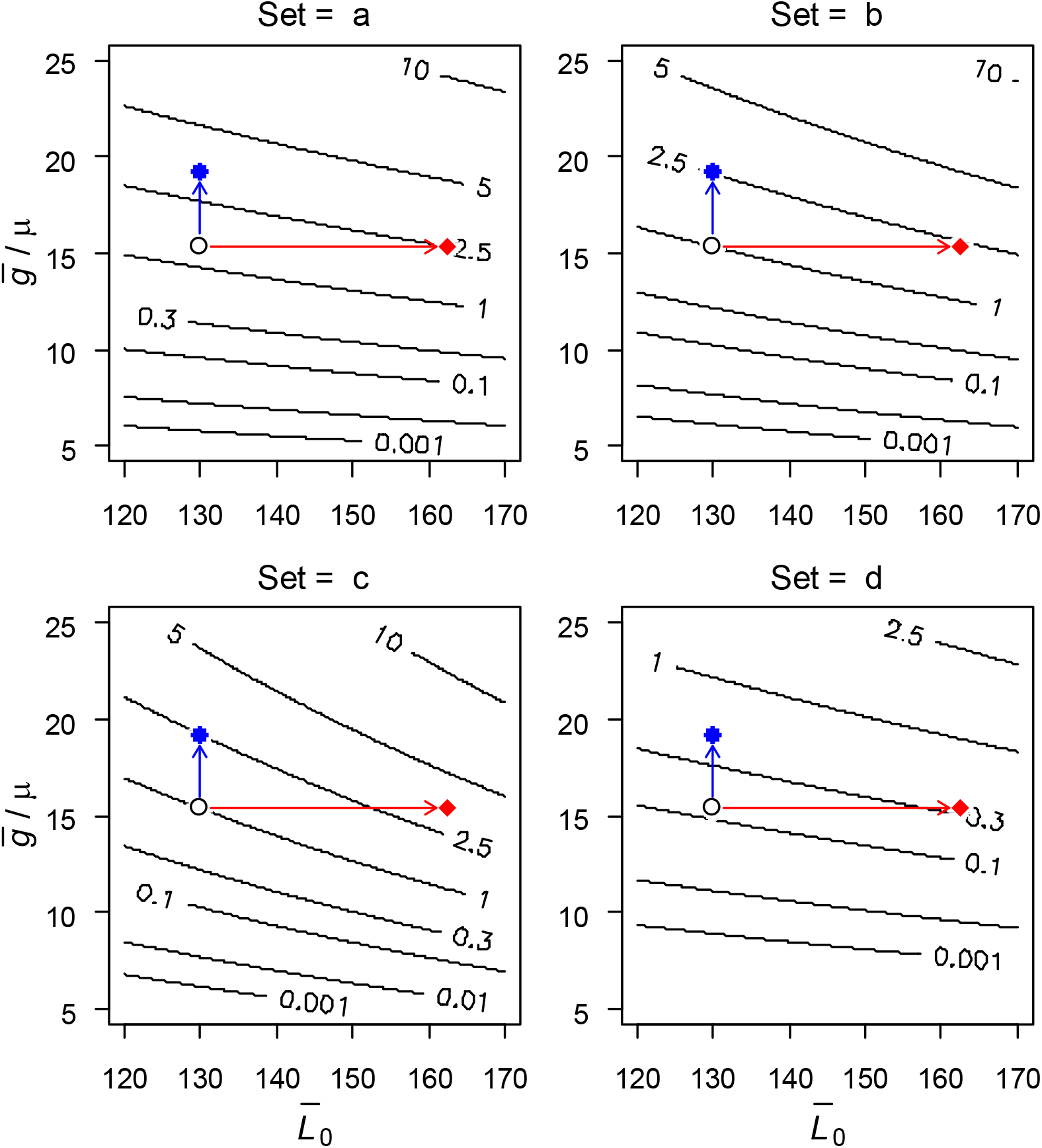
Contours of SAR (%) as a function of juvenile fish length at ocean entrance vs the ratio of the rates of growth to mortality for predator distribution sets a-d in table 1. Based conditions (∘) *ḡ/μ* and 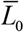 are same in each set. Parameters increases of 25% are depicted as 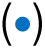 for *ḡ/μ* and 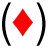 for 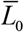.

## 4. Discussion

The model describes survival patterns of prey fish exposed to gape-limited predators in terms of the amount of time it takes the prey to grow larger than the predator gapes. The approach has a Lagrangian framework and expresses the probability of prey mortality in terms of the frequency of predator-prey encounters and the relative size differential at the encounters. While a prey-centric perspective has been applied in survival models [e.g. 27] the model here is unique in coupling the predator size distribution with the size and growth of the prey.

The model provides a mechanistic explanation for statistical analyses that demonstrate juvenile fish size and growth correlate with adult survival and recruitment. The properties of this framework are illustrated using Chinook salmon from the Snake River in the Columbia Basin of the Pacific Northwest. The estimated model parameters favorably compare to independent estimates of early marine growth and mortality in this population. While commercially managed stocks are seldom monitored at the level of the Snake River salmon, which are endangered, it is plausible that even limited size information from commercial stocks could be used to improve recruitment predictions. For example, measures of size of migrating juveniles and early maturing spawners may be sufficient to estimate the growth rate parameters which could then be used to predict ocean survival of later returning spawners. Additionally, otolith-based estimates of early marine growth [31] would be especially informative for predicting fish returns.

A central goal in fisheries management is to understand the effects of growth and size on survival and recruitment. Theories have suggested fish need to reach a critical size by a particular life stage [33, 34] while others have demonstrated the importance of early marine growth rates [31, 35, 36]. Furthermore, other studies suggest freshwater growth prior to migration affect adult survival [6, 12, 37, 38]. The properties noted in these studies and others are intrinsic to the model. In addition, the model includes the heretofore unconsidered mechanism by which the predator size distribution affects survival.

The model’s Lagrangian framework may also provide a tractable approach for linking prey and predator dynamics in ecological theory. Traditional predator-prey theory cast mortality in terms of predator and prey densities and encounter frequencies but is largely silent on the age-dependent effects of the interactions. The model presented here, by linking relative predator size and prey growth opens a potentially new perspective. For example, the strength of density-dependent mortality would depend on the effect of prey density on prey growth rate as well as the size-frequency distribution of predator gapes. Consequently, the strength of density dependence could be inversely correlated with the variance in the predator size-frequency distribution and directly correlated with the strength of prey competition for food resource. In a similar manner, the predator size frequency distribution and prey growth rate may play a role in food web stability. For example, the persistence of a prey trophic level might be expected to depend on the prey individual growth rate and the size frequency distribution of the associated predator trophic level. In essence, to avoid extirpation the prey’s growth would need to be sufficient to allow a critical number of the population to outgrow the size frequency distribution of the predators. In a final example, the model framework might be suitable to explore carryover effects in which early life conditions affect survival in later life stages [39]. For example, the SAR of hatchery reared Chinook salmon was found to be correlated with the environmental conditions prior to juvenile migration. The mechanism is thought to involve the physiological development of growth factors and smoltification prior to juvenile migration to the ocean [12]. These mechanisms might be readily incorporated into the model, and therefore into adult survival projections by linking the marine growth rate parameter to the early life physiological development.

In summary, over several decades of research, fish growth and size have emerged as central factors determining population survival and recruitment. This paper expresses mortality in terms of prey growing through a field of predator gapes to show that the recruitment strength can depend on the size distribution of the predators. Studies exploring the strength and significance of this linkage will be required to assess its importance to predator-prey theory and the management of commercially important fishes.

## Data accessibility

Data available in PITAGS

## Authors’ contributions

JA is sole author

## Acknowledgements

I wish to thank Jeff Rudder for helpful comments on the model development and Jennifer Gosselin for valuable editorial comments

